# “Δ133p53α Protects Human Astrocytes from Amyloid-Beta Induced Senescence and Neurotoxicity”

**DOI:** 10.1101/2021.05.17.444544

**Authors:** Kyra Ungerleider, Delphine Lissa, Jessica A. Beck, Izumi Horikawa, Curtis C. Harris

## Abstract

Cellular senescence is an important contributor to aging and age-related diseases such as Alzheimer’s disease (AD). Senescent cells are characterized by a durable cell proliferation arrest and the acquisition of a proinflammatory senescence-associated secretory phenotype (SASP), which participates in the progression of neurodegenerative disorders. Clearance of senescent glial cells in an AD mouse model prevented cognitive decline suggesting pharmacological agents targeting cellular senescence might provide novel therapeutic approaches for AD. Δ133p53α, a natural protein isoform of p53, was previously shown to be a negative regulator of cellular senescence in primary human astrocyte, with clinical implications from its diminished expression in brain tissues from AD patients. Here we show that treatment of proliferating human astrocytes with amyloid-beta oligomers (Aβ), an endogenous pathogenic agent of AD, results in reduced expression of Δ133p53α, as well as induces the cells to become senescent and express proinflammatory SASP cytokines such as IL-6, IL-1β and TNFα. Our data suggest that Aβ-induced astrocyte cellular senescence is associated with accelerated DNA damage, and upregulation of full-length p53 and its senescence-inducing target gene p21^WAF1^. We also show that exogenously enhanced expression of Δ133p53α rescues human astrocytes from Aβ-induced cellular senescence and SASP through both protection from DNA damage and dominant-negative inhibition of full-length p53, leading to inhibition of Aβ-induced, astrocyte-mediated neurotoxicity. The results presented here demonstrate that Δ133p53α manipulation could modulate cellular senescence in the context of AD, possibly opening new therapeutic avenues.

**HIGHLIGHTS:**

- Aβ diminishes the expression of the p53 isoform Δ133p53α, and induces cellular senescence, SASP, and DNA damage in human astrocytes.
- Δ133p53α protects human astrocytes from Aβ-induced DNA damage and cellular senescence.
- Δ133p53α prevents astrocyte-mediated neurotoxicity to neuro-progenitor cells.

## INTRODUCTION

There is a critical need to identify novel therapeutic targets in the treatment of age-associated diseases including dementia. Currently, 5.7 million Americans are living with Alzheimer’s disease (AD) and by 2050, this number is projected to reach nearly 14 million [1].

AD is characterized by the accumulation of amyloid-beta (Aβ) plaques produced from amyloid precursor protein (APP). APP is a transmembrane protein cleaved by β-site APP cleaving enzyme 1 (known as BACE1) and γ-secretase to produce Aβ fragments between 37-43 amino acids long [2]. These Aβ peptides aggregate and contribute to neurotoxicity and subsequent neurocognitive decline [3]. The accumulation of Aβ increases with normal aging [4], and in a more accelerated manner in AD. In association with Aβ accumulations in the brain, several brain cell types, including astrocytes and microglia, adopt an inflammatory phenotype and function to clear Aβ from the extracellular space [5]. However, prolonged activation of these glial cells in the presence of Aβ contributes to the chronic inflammation observed in AD. Although several forms of Aβ exist, there is increased interest in the soluble species of Aβ, particularly the oligomeric form such as Aβ_1-42_, which correlates better with disease severity [6].

Cellular senescence is characterized by sustained cell cycle arrest and is induced by telomere shortening (referred to as replicative senescence) or by cellular stressors inducing DNA damage and activating oncogenes. Common markers of cellular senescence include upregulation of cell cycle inhibitors p16^INK4A^ [7] and p21^WAF1^ [8], as well as increased activity of senescence-associated β-galactosidase (SA-β-gal), a lysosomal hydrolytic enzyme [9]. When cells undergo senescence, they secrete various inflammatory cytokines which alter the tissue microenvironment and propagate neurodegeneration [10]. This inflammatory phenotype is collectively named the senescence-associated secretory phenotype, or SASP, but may vary depending on the cell type and the inducer of senescence [11]. Astrocytes, an important regulator of brain homeostasis providing functional and metabolic support [12], are a major cell type undergoing senescence in the brain [13]. Senescent astrocytes, which are induced by aging- or injury-associated replicative exhaustion [13] or in response to endogenous and external stresses such as irradiation [14] show phenotypic similarities to a subset of so-called reactive astrocytes in terms of increased secretion of pro-inflammatory SASP cytokines and neurotoxicity [15]. Here, we show for the first time that Aβ induces astrocytic SASP, and identify a role for astrocytic SASP in Aβ promoted neurotoxicity. Aβ oligomers have been reported to induce senescence in astrocytes [16] but the mechanism underlying Aβ-induced senescence is still to be elucidated.

TP53 plays a critical role in the initiation of cellular senescence through upregulating p53-inducible senescence genes such as p21^WAF1^ [17]. Elevated p53 immunoreactivity has been observed in sporadic and familial AD, particularly in white matter glial cells distributed in brain regions undergoing degeneration [18], suggesting that p53 and its regulatory factors in glial cells are involved in AD pathogenesis. The human TP53 gene encodes, in addition to full-length p53 protein (often simply called p53), at least 12 natural protein isoforms through alternative mRNA splicing or alternative promoter usage [19]. Δ133p53α, an N-terminal truncated isoform, dominant-negatively inhibits full-length p53-mediated senescence and promotes DNA repair under conditions of accelerating DNA damage [14, 20–22]. Expression of Δ133p53α was diminished in the brain tissue of AD patients, compared to age-matched controls [13]. We have shown astrocytes to be one of the predominant cell types expressing Δ133p53α in the brain [13, 14]. Decreased expression of Δ133p53α was associated with radiation-induced and replicative senescence in human astrocytes, which was rescued by overexpression of Δ133p53 [13, 14]. All these findings have prompted us to investigate whether Δ133p53α plays a role in the underlying mechanism for Aβ-induced astrocyte senescence and SASP. It is unknown whether AD pathogenic agents such as Aβ directly induce this astrocyte phenotype and whether Δ133p53α has a protective effect against Aβ-induced astrocytic neurotoxicity. In this study, we show that treatment of primary human astrocytes with Aβ causes reduced expression of Δ133p53α and accelerated DNA damage, along with induction of senescence and SASP, and that lentiviral vector-based expression of Δ133p53α promotes DNA repair in Aβ-treated astrocytes and prevents them from SASP-mediated neurotoxicity. Together with a therapeutic effect of clearance of senescent glial cells against cognitive decline in mouse models [23, 24], this study using human astrocytes emphasizes a causative role of astrocyte senescence in neurodegenerative diseases and supports Δ133p53α as a novel therapeutic target that can be leveraged in astrocytes.

## EXPERIMENTAL PROCEDURE

### Primary cells and cell lines

Primary human astrocytes were purchased from Sciencell (Carlsbad, CA, USA) and cultured at 37°C, 5% CO_2_ in astrocyte medium supplemented with 2% fetal bovine serum, growth supplement, and 1% penicillin/streptomycin all obtained from ScienCell Research Laboratories (Carlsbad, CA). Cells were passaged when confluent. Human neural progenitor cells (NPCs) (ACS-5004) were maintained in neural expansion media (ACS-3003) purchased from American Type Tissue Collection (Manassass, VA) passaged when 80% confluent.

### Preparation of Oligomeric Amyloid-beta

Synthetic Aβ_1-42_ pretreated with hexafluoroisopropanol HFIP (1,1,1,3,3,3-hexafluoro-2-propanol) was purchased from Bachem (Switzerland) and oligomerized as described [25]. Briefly, to generate oligomeric Aβ, the peptide film was dissolved in DMSO to a concentration of 5 mM and sonicated. The suspension was diluted in phenol-free DMEM to a final concentration of 100 μM and incubated at 4°C overnight.

### Cell Treatments

Human astrocytes were treated for 24 hours with oligomerized Aβ_1-42_ at a final concentration of 0.5 or 1 μM (as indicated in Figure legends) in astrocyte media; 0.01% DMSO was used as control. After 24 hours, media was replaced without Aβ. Cell lysates were collected 72 hours days post-treatment for immunoblotting, and qRT-PCR.

### Lentiviral vector transduction

As described previously [13, 21], the pLOC lentiviral vector, which drives RFP as an ORF insert and IRES-translated GFP, was purchased from Open Biosystems (Lafayette, CO). For Δ133p53α expression, RFP was replaced with the Δ133p53α cDNA. These lentiviral constructs, together with the Trans-Lentiviral GIPZ packaging system (Open Biosystem), were transfected into 293T/17 Cells (ATCC, American Type Culture Collection) using Lipofectamine-2000 (Invitrogen). The Δ133p53 cDNA was also cloned into pLenti6.3/TO/V5-DEST vector (Thermo Fisher Scientific) by using Spe I and Mlu I sites for constitutive overexpression and were transiently transfected into 293T/17 cells (ATCC) with the ViraPower lentiviral expression system (Thermo Fisher Scientific). The lentiviral vector particles were collected 48 hours after transfection and used for overnight transduction to primary human astrocytes, followed by selection with blasticidin (1 μg/ml; Sigma–Aldrich) three days after transduction. The pLenti6.3/TO/V5-DEST lentiviral vector was used for Fig. 3 and the RFP/GFP pLOC lentiviral vector was used for all other experiments.

**Fig. 1:**
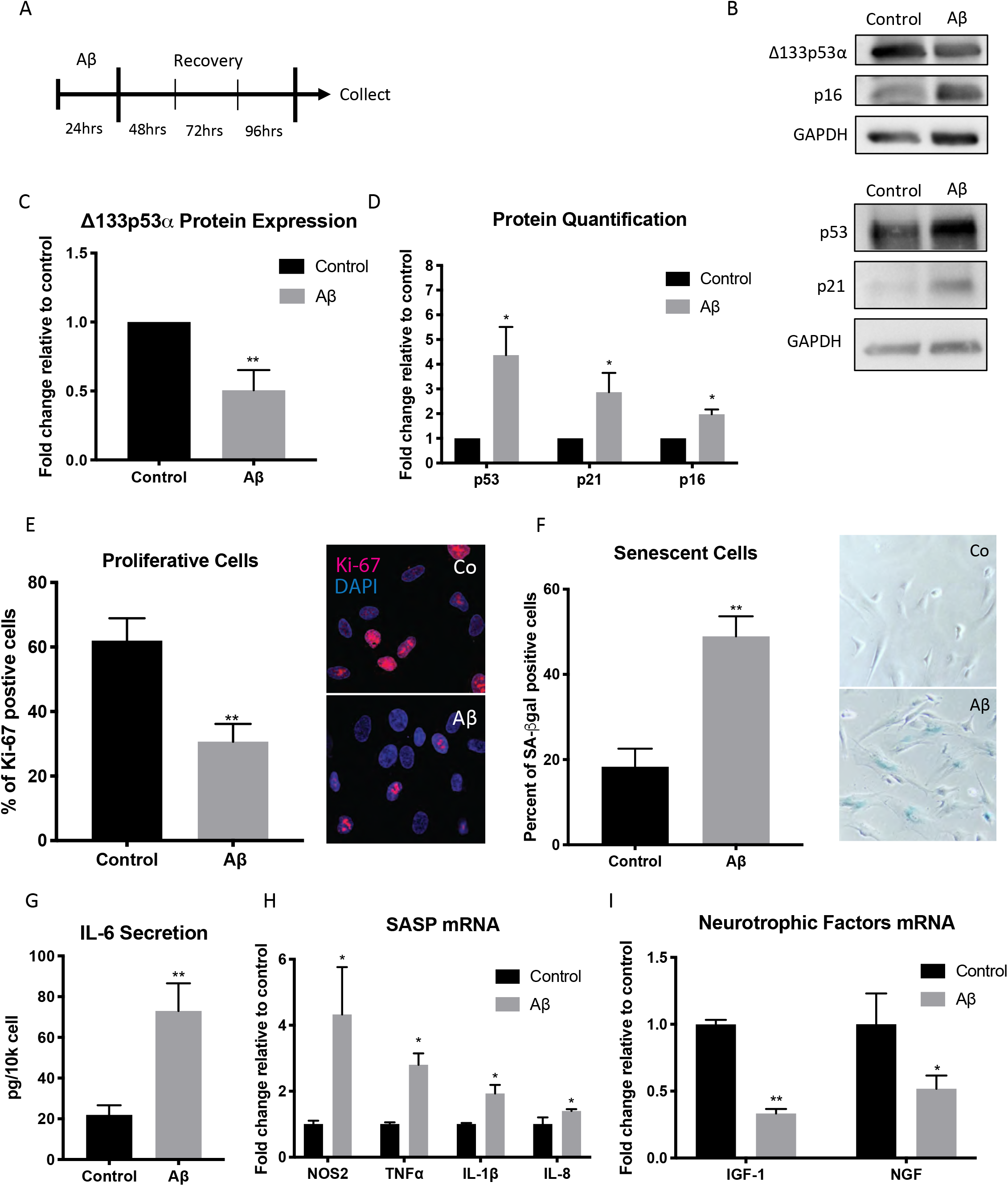
Amyloid-beta oligomers induce senescence and SASP in human astrocytes. **(A)** Timeline of experiment. Human astrocytes were treated for 24 hours at 0.5 μM or 1 μM Aβ. Media was changed after 24 hours and cells were collected or stained 72 hours after treatment end. Control astrocytes were incubated with 0.01% DMSO, instead of Aβ. **(B)** Representative immunoblot images of astrocytes treated with 0.5 μM Aβ as in (A). GAPDH was for normalization. **(C, D)** Quantitative analysis of protein expression levels of Δ133p53α (C) and p53, p21^WAF1^ and p16^INK4A^ (D). The expression levels (normalized to GAPDH) in Aβ-treated cells are shown relative to those in Control. (**E)** Immunofluorescence staining images (40x magnification) and quantitative analysis of proliferation marker Ki-67 (red), with DAPI nuclear staining (blue), in astrocytes treated with 1 μM Aβ. **(F)** Representative images of SA-β-gal staining (20x magnification) and quantification of percent positive (blue) cells in control and Aβ-treated astrocytes (0.5 μM). **(G)** Quantification of IL-6 in astrocyte media following 72-hr recovery from 0.5 μM Aβ treatment, as shown in (A). The data were normalized to the cell numbers (picogram per 1 × 10^4^ cells) **(H, I)** Fold change of mRNA expression of NOS2, TNFα, IL-1β and IL-8 (H), and of IGF-1 and NGF (I) in astrocytes treated with 0.5 μM Aβ. The expression levels were normalized to GAPDH. Data are presented as mean ± S.E.M. * *p*≤0.05, \*\**p* ≤0.001.

**Fig. 2:**
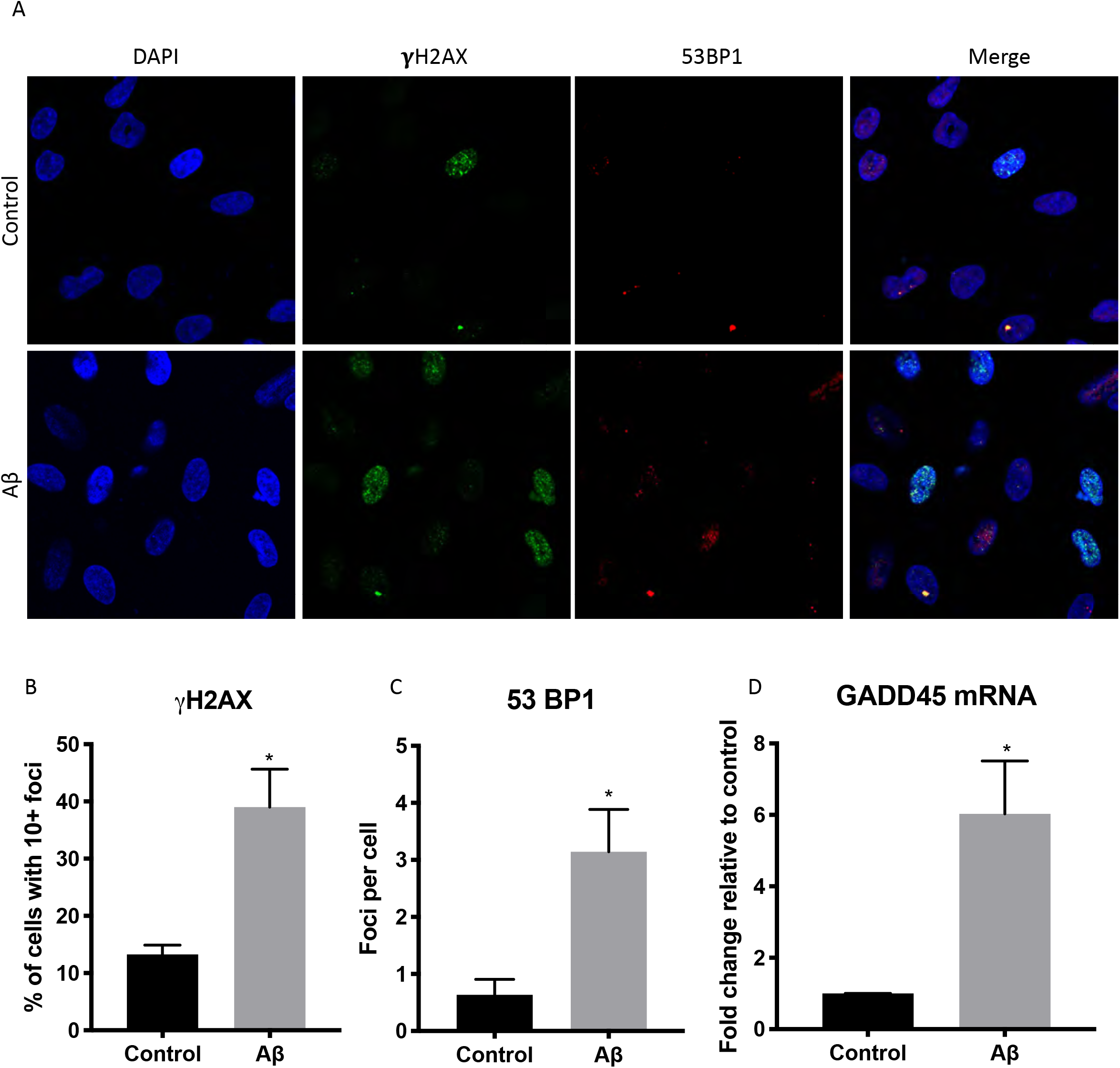
Amyloid-beta oligomers induce DNA damage in human astrocytes. **(A)** Representative images of immunofluorescence labeling of double-stranded DNA breaks by γH2AX (green) and 53BP1 (red) 5 days after treatment with 1 μM Aβ for 24 hours (40x magnification). **(B, C)** Quantitative analysis of DNA damage foci of γH2AX (B) and 53BP1 (C). **(D)** Fold change of GADD45 mRNA expression normalized to GAPDH. Data are presented as mean ± S.E.M. \**p*≤0.05.

**Fig. 3:**
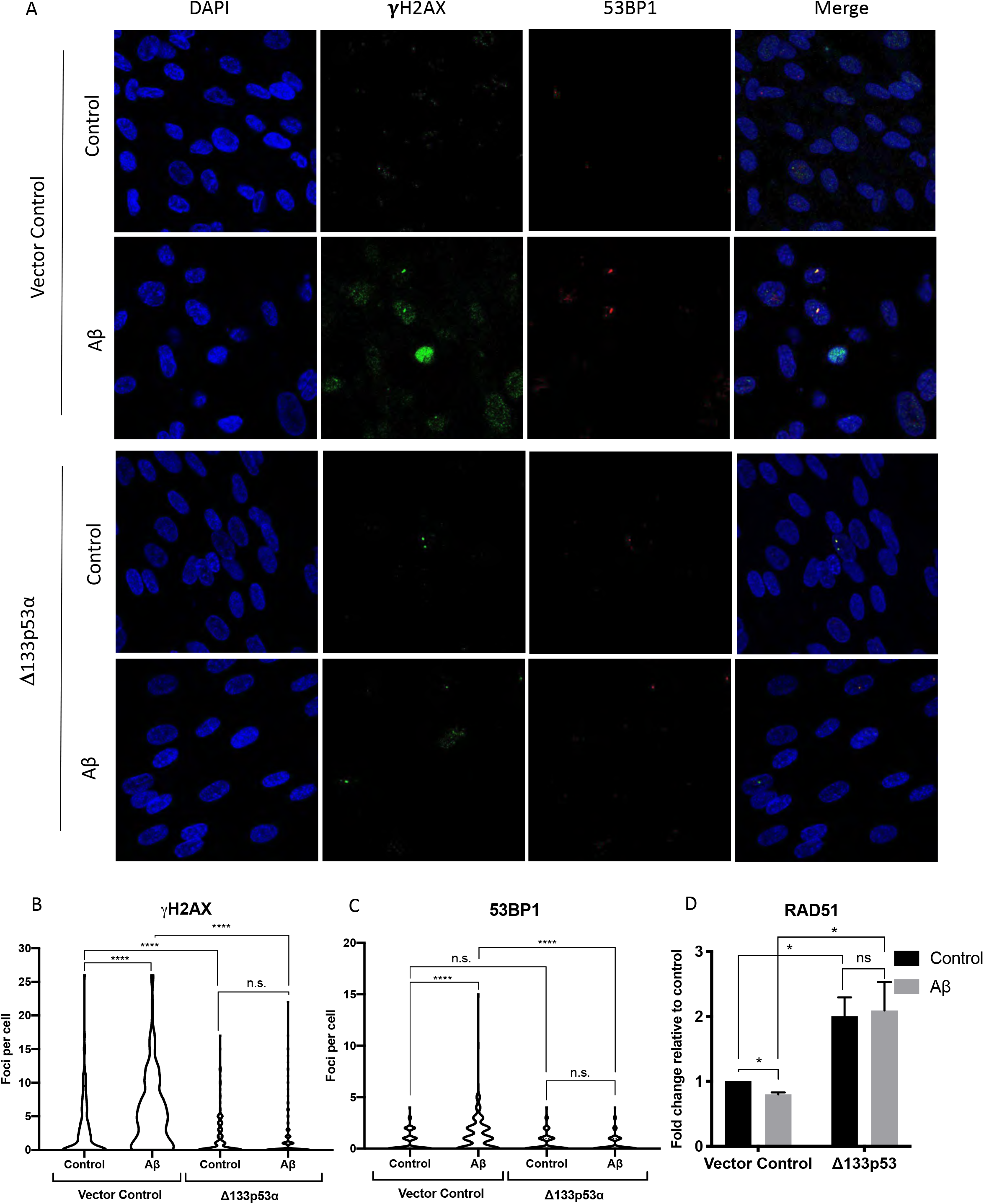
Δ133p53α protects from amyloid-beta induced DNA damage. **(A)** Representative images of immunofluorescence staining of double-stranded DNA breaks by γH2AX (green) and 53BP1 (red). Vector control and Δ133p53α-expressing astrocytes were treated with Aβ (or control) and recovered, as in Fig 2. (40x magnification). **(B, C)** Quantification of numbers of γH2AX foci (**B**) and 53BP1 foci (**C**) per cell. The pooled data from three separate experiments are shown as a single graph for each group. **(D)** Fold change of RAD51 mRNA expression normalized to GAPDH. Data are presented as mean ± S.E.M. \**p*≤0.05. \*\*\*\**p*≤0.0001, n.s. = not significant.

### Trans-well Experiments

Human astrocytes were treated with Aβ_1-42_ in the top transwell chamber for 24 hours then the media was changed without Aβ_1-42_ and combined with NPCs. NPCs were plated on bottom transwell chamber 24 hours before combining with human astrocytes in 1:1 astrocyte and NPC media. After 72hr co-culture, accutase was used to collect NPCs for trypan blue staining and qRT-PCR.

### SA-β-gal staining

SA-β-gal staining was performed according to the protocol of the Senescence-β-Galactosidase Staining Kit (Cell Signaling Technology, Danvers, MA, USA). At least 200 cells were counted in each treatment group from three independent experiments.

### IL-6 Quantification

Quantification of IL-6 in media was assessed via Human IL-6 ELISA kit according to manufacturer instructions (Sigma Aldrich; St. Louis, MO).

### Immunofluorescence

Cells were plated onto poly-D-lysine/laminin coated 8-chamber slides (Corning). After 24 hours, cells were washed with PBS and fixed for 10 min with 4% paraformaldehyde. Cells were permeabilized with 0.01% Triton-X for 4 min, washed with PBS and then blocked in 1% BSA for 1 h at room temperature. Primary antibodies listed in Supplementary Table 1 were applied overnight at 4 °C. Cells were washed with PBS before incubation with a secondary antibody conjugated to fluorophores: Alexa-488, 568- and −647 at a dilution of 1:400 (Life Technologies). Coverslips were mounted on with Vectashield Mounting Medium with DAPI (Vector Laboratories, Inc.; Burlingame, CA).

### Immunoblotting

Immunoblotting was carried out as previously described [14]. Briefly, cells were lysed in radioimmunoprecipitation assay (RIPA) buffer and protein concentration was determined by Bradford assay. NuPAGE 4 × loading buffer was added to all lysates and then boiled for 5 min before loading onto a Tris-glycine gel (Thermo Scientific, Waltham, MA, USA) for electrophoresis. Proteins were then transferred onto a PVDF membrane. Membranes were blocked in 1:1 mixture of Superblock and Tris-Buffered Saline (TBS, 125 mM Tris and 200 mM NaCl), containing 0.1% Tween-20 and 10% non-fat dry milk. Membranes were incubated in the primary antibodies listed in Supplementary Table 1 overnight at 4 °C, and washed three times in TBS-Tween-20. Membranes were then incubated in HRP-conjugated secondary antibody for 1 h at room temperature. The signal was visualized with SuperSignal developing reagent (Thermo Fisher Scientific) using ChemiDoc Touch Imaging System (BioRad Laboratories) and ImageLab software was used for quantitative image data analysis.

### RNA extraction and cDNA preparation

Total RNA was extracted using the RNeasy Mini Kit (Qiagen, Crawley, UK) according to the manufacturer’s instructions. Cells were homogenized and lysate mixed 1:1 with 70% ethanol and centrifuged through an RNeasy Mini Spin column. RNA was eluted with RNase-free water. The abundance and quality of the resulting RNA was assessed using a Nanodrop ND-1000 spectrophotometer (Nanodrop Technologies, Wilmington, DE, USA). RNA samples were diluted so that 2000 ng total RNA could be used for a 5-μl reverse-transcription reaction. cDNA was synthesized using SuperScript II Reverse Transcriptase (Invitrogen).

### Quantitative real-time polymerase chain reaction (qRT-PCR)

For the quantitative analysis of mRNA expression, the CFX384 Real-Time PCR system (BioRad Laboratories Hercules, CA, USA) was used. Each reaction was performed in at least triplicate using 2 μl of cDNA in a final volume of 5 μl. The following thermal cycle was used for all samples: 10 min-95 °C; 40 cycles of 30 s-95 °C, 40 s-primer specific annealing temperatures, 40 s-72 °C. The expression level of each target gene was analyzed based on the ΔΔCt method and the results expressed as relative expression normalized to GAPDH. Taqman primer assays for p21 (Hs00355782_m1), NOS2 (Hs01075529_m1), IL-8 (Hs00174103_m1), IL-1β (Hs01555410_m1), TNFα (Hs00174128_m1), GADD45 (Hs00169255_m1), BAX (Hs00180269_m1), PUMA (Hs00248075_m1), NGF (Hs00171458_m1), IFG1 (Hs01547656_m1), and GAPDH (Hs02758991_g1) were purchased from Life Technologies (Thermo Fisher Scientific).

### Statistics

Data are presented as mean ± S.E.M. of at least 3 independent experiments. Comparisons were made using 2-sided, paired Student’s *t*-test for all qRT-PCRs and immunoblotting. All other assays were analyzed by using 2-sided, unpaired Student’s *t*-test. Differences were considered significant at \**p* ≤ 0.05, **p ≤ 0.01, ***p ≤ 0.001, \*\*\*\**p*≤0.0001.

## RESULTS

### Aβ Decreases Expression of Δ133p53α and Induces Senescence in Human Astrocytes

In order to assess Aβ effect on astrocyte senescence and p53 isoform expression, primary human astrocytes were treated with oligomeric Aβ_1-42_ (referred to as Aβ for short from here out) for 24 hours then collected after a 3 day “recovery period” in fresh media (Fig. 1A). Treatment with Aβ increased expression of p53, its target cell cycle inhibitor p21^WAF1^ and another cell cycle inhibitor p16^INK4A^, while decreasing endogenous Δ133p53α expression (Fig. 1B-D). Aβ inhibited expression of cell proliferation marker Ki-67 (Fig. 1E), and increased SA-β-gal activity (Fig. 1F), a marker of cellular senescence. These results indicate that Aβ leads to activation of p53, with downregulation of its inhibitory isoform Δ133p53α, and induces senescence in human astrocytes.

### Astrocytes Display SASP Following Aβ Treatment

IL-6 is a major contributor to SASP and is often increased in aging-related diseases [26]. Astrocytes treated with Aβ demonstrated increased IL-6 secretion (Fig. 1G) and elevated mRNA expression of common SASP cytokines including TNFα, IL-1β and IL-8, and of NOS2 (iNOS) associated with SASP and oxidative stress (Fig. 1H). Additionally, astrocytes have reduced expression of neurotrophic factors (IGF-1 and NGF) (Fig. 1I). These results further support the induction of senescence by Aβ and identify features of proinflammatory and neurotoxic SASP in Aβ-treated human astrocytes.

### Aβ Induces DNA Damage in Astrocytes

DNA damage contributes to the development of AD and has been observed in astrocytes found in AD vulnerable regions of the brain [27–29]. Although DNA damage-induced transient cell cycle arrest is a mechanism for preventing the replication of cells with potential oncogenic mutations [30], increased accumulation of DNA damage in persistent senescent cells can contribute to aging-associated diseases. In the case of Aβ, Aβ_1-42_ peptide oligomers demonstrate DNA nicking activities *in vitro* similar to nucleases [31] and Aβ-associated oxidative stress could lead to DNA double-stranded breaks [32]. To examine if Aβ-induced astrocyte senescence is associated with DNA damage, astrocytes were treated with Aβ_1-42_ as above and labeled for double-stranded DNA damage markers γH2AX and 53BP1. We observed increased numbers of γH2AX- and 53BP1-positive foci in astrocytes treated with to Aβ (Fig. 2A-C, Supplementary Fig. 1A and B), as well as elevated mRNA expression of GADD45, a DNA damage-inducible gene (Fig. 2D). These results indicate that Aβ induces DNA damage in astrocytes, which could consequently contribute to the induction of senescence.

### Δ133p53α Inhibits Aβ-induced DNA Damage

Previous studies have shown that Δ133p53α promotes DNA repair in fibroblasts and cancer cells [22, 33]. To examine whether Δ133p53α also protects human astrocytes from Aβ-accelerated DNA damage, we generated human astrocytes expressing the lentiviral vector-driven Δ133p53α (Supplementary Fig. 2). Although Δ133p53α reduced baseline levels of γH2AX foci in control cells without Aβ, the more striking effect of Δ133p53α was observed against Aβ-induced DNA damage (Fig. 3A and B). The Δ133p53α-expressing astrocytes, whether or not treated with Aβ, showed only modest numbers of γH2AX foci, suggesting that they were completely resistant to Aβ-induced DNA damage (Fig. 3B). The protective effect of Δ133p53α against Aβ-induced DNA damage was also confirmed by the analysis of 53BP1 foci (Fig. 3A and C). Consistent with previous findings [22], RAD51 was upregulated by Δ133p53α in the presence or absence of Aβ (Fig. 3D), suggesting that the effect of Δ133p53α may be attributed in part to increased expression of this DNA repair protein.

### Δ133p53α Protects Astrocytes from Aβ-induced Senescence

To determine whether the lentiviral vector-driven expression of Δ133p53α would protect from Aβ-induced senescence, control vector-transduced and Δ133p53α-expressing primary human astrocytes were examined for senescent phenotypes. We found that Δ133p53α-expressing astrocytes showed no significant increase in SA-β-gal staining after Aβ treatment, compared to an increase up to 50% positivity in control astrocytes (Fig. 4A, B). The Δ133p53α-expressing astrocytes expressed lower levels of p21 mRNA, which did not respond to Aβ, in contrast to the control cells showing the higher basal level and its further increase in response to Aβ (Fig. 4C). Similarly, the Δ133p53α-expressing astrocytes secreted less IL-6 and there was no induction by Aβ treatment, while Aβ significantly increased the IL-6 secretion in the control cells (Fig. 4D). Aβ treatment also induced the TNFα expression in the control cells, but not in the Δ133p53α-expressing astrocytes (Fig. 4E). In addition, the lentiviral expression of Δ133p53α upregulated neurotrophic IGF-1 and NGF and rendered them resistant to reduction by Aβ (Fig. 4F, G). These results indicate that Δ133p53α protects astrocytes from Aβ-induced senescence and SASP, along with upregulation of the neurotrophic growth factors.

**Fig. 4:**
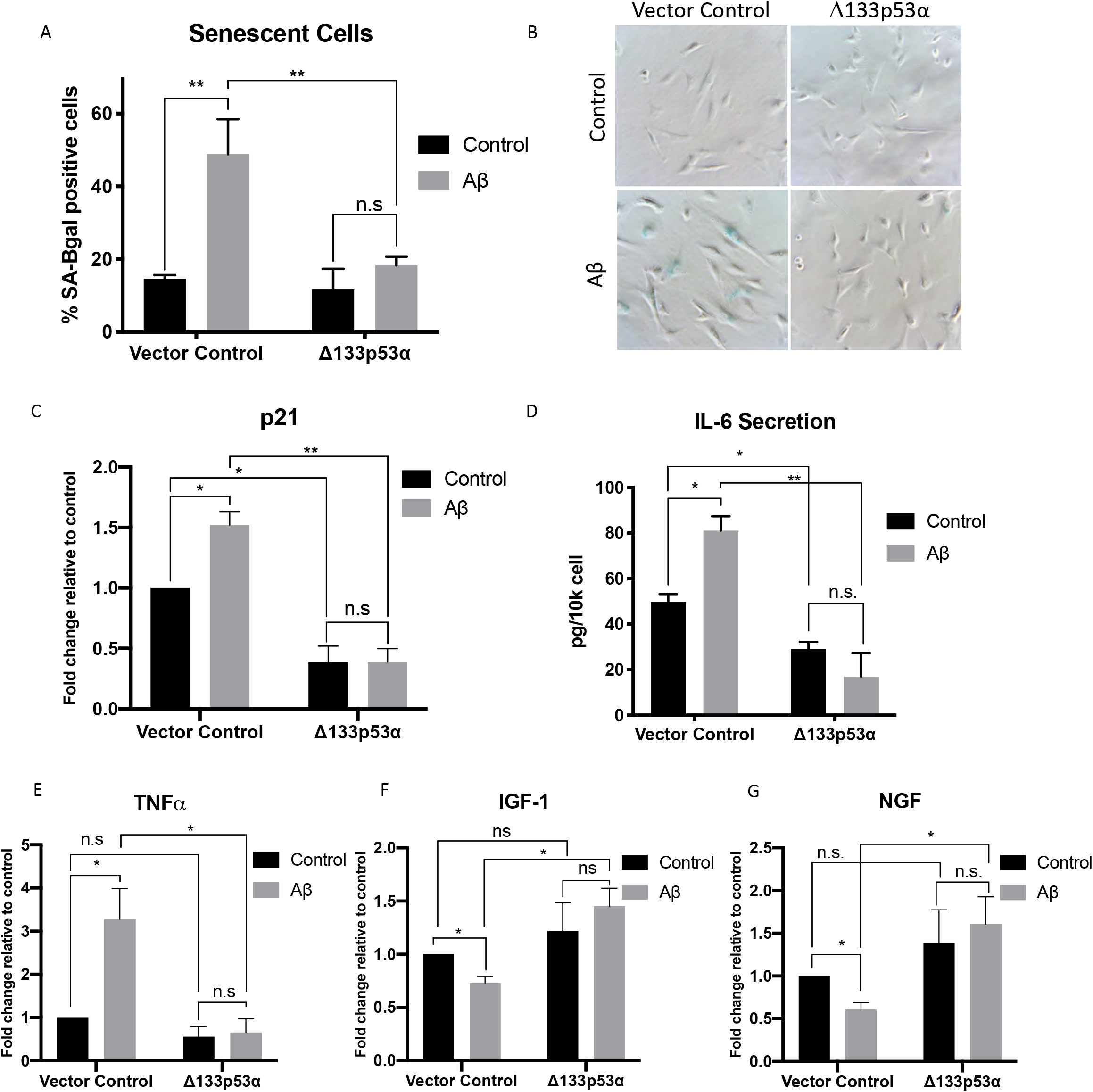
Δ133p53α inhibits amyloid-beta induced astrocyte senescence. Astrocytes expressing the lentiviral vector-driven Δ133p53α and with the vector control were treated (for 24 hours) with 0.5 μM Aβ and recovered (for 72 hours), along with control without Aβ treatment, as shown in Fig. 1A. **(A, B)** SA-β-gal staining. Quantification of positive cells (A) and representative images (10x magnification) (B). **(C)** Fold change of p21 mRNA expression normalized to GAPDH. **(D)** Quantification of IL-6 secretion in astrocyte media, as in Fig. 1G. **(E-G)** Fold change of mRNA expression of TNFα (E), IGF-1 (F) and NGF (G). The expression levels were normalized to GAPDH. Data are presented as mean ± S.E.M. \**p*≤0.05. \*\**p*≤0.01, n.s. = not significant.

### Δ133p53α Prevents Astrocyte-mediated Neurotoxicity Following Aβ Treatment

Astrocyte dysfunction can result in NPC death, further propagating neurodegeneration [12]. To investigate the protective role of Δ133p53α in Aβ-induced astrocyte neurotoxicity, we cocultured transduced astrocytes expressing Δ133p53α and control vector with NPCs separated by a trans-well membrane. Astrocytes were treated with Aβ for 24 hours, then fed with fresh media without Aβ, and the astrocytes were combined with NPCs in trans-well system. NPCs were in co-culture with astrocytes for 3 days and then analyzed or collected (Fig. 5A). Cell viability measured by trypan blue was decreased in NPCs co-cultured with vector control astrocytes exposed to Aβ, but no significant loss of cell viability was observed in NPCs co-cultured with Δl33p53α-expressing, Aβ-exposed astrocytes (Fig. 5B). To determine if the NPCs were undergoing apoptosis, we quantified cleaved caspase 3 positivity via immunofluorescence and observed an increase of cleaved caspase 3-positive cells in NPCs co-cultured with Aβ-treated vector control astrocytes, but not in NPCs co-cultured with astrocytes expressing Δ133p53α with or without Aβ treatment (Fig. 5C, D). Additionally, we found increased expression of proapoptotic genes BAX and PUMA and a cell cycle inhibitor p21 in NPCs co-cultured with Aβ-exposed vector control astrocytes, but again not in those co-cultured with astrocytes expressing Δ133p53α (Fig. 5E-G). These findings indicate that Δ133p53α expression in astrocytes inhibits Aβ-induced, astrocytic SASP-mediated neurotoxicity.

**Fig. 5:**
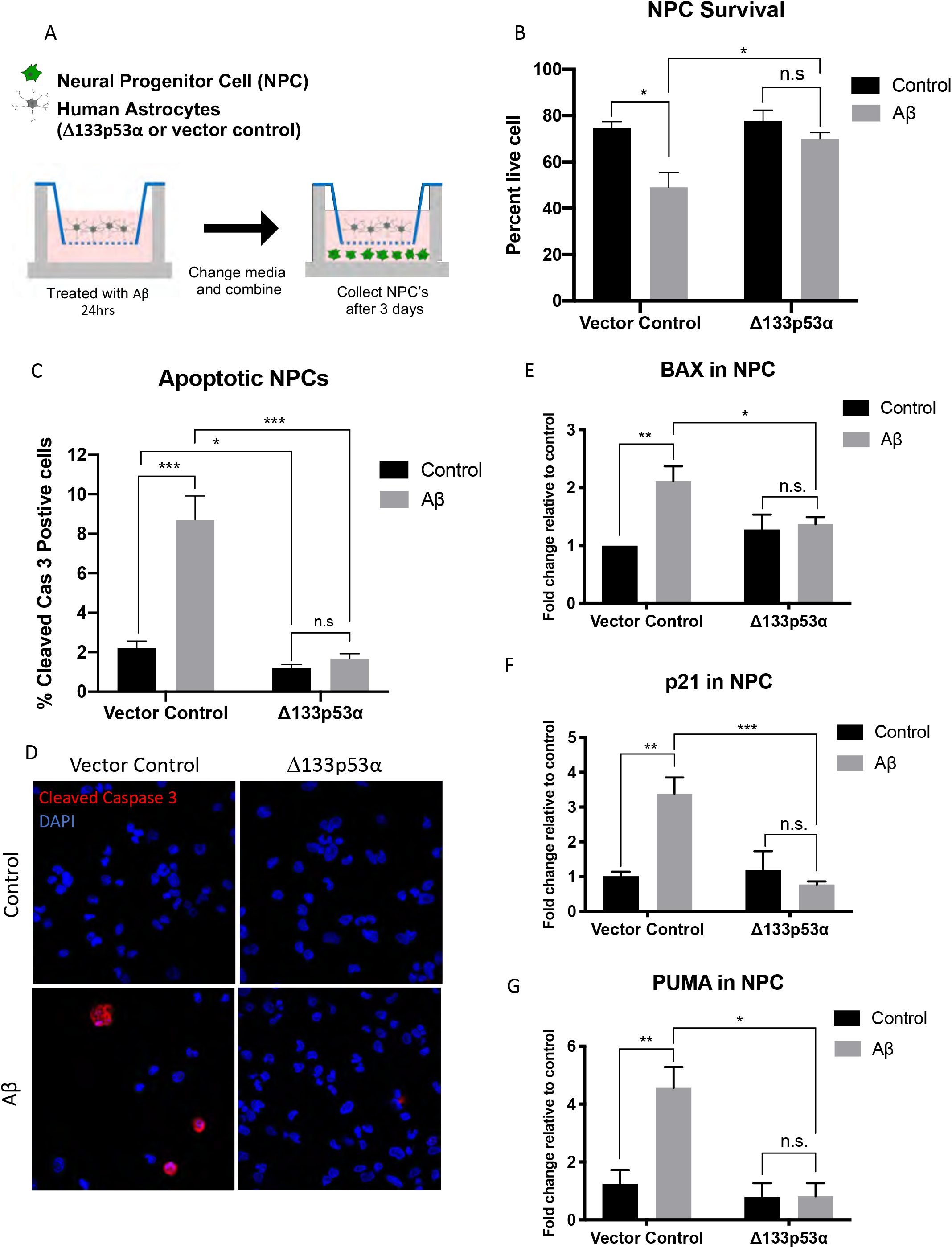
Δ133p53α inhibits amyloid-beta induced astrocyte neurotoxicity to Neuroprogenitor Cells. Astrocytes with the Δ133p53α expression and with the vector control were treated for 24 hours with 0.5 μM Aβ in top of trans-well. Media was then changed without Aβ and astrocytes were combined with NPCs plated on bottom of trans-well. NPCs were analyzed after 72-hour co-culture with astrocytes. **(A)** Illustration of experiment design and timeline. **(B)** Cell survival measured by trypan blue exclusion. **(C, D)** Immunofluorescence staining of cleaved caspase 3 in apoptotic NPCs. Quantitative data (C) and representative images (40x magnification) (D) are shown. **(E-G)** Fold change of mRNA expression of BAX (E), p21 (F) and PUMA (G) in NPCs. The expression levels were normalized to GAPDH. Data are presented as mean ± S.E.M. \**p*≤0.05, \*\**p*≤0.01. \*\*\**p*≤0.001, n.s. = not significant.

## DISCUSSION

Astrocytes provide neurotrophic support to neurons [5], however with physiological and accelerated aging, endogenous and exogenous pathogenic agents and stimuli, and chronic stress and injury, they may become dysregulated and dysfunctional [15]. Aβ represents one such endogenous agent for AD pathogenesis, which may lead to chronic oxidative stress and aging-associated phenotypes [28], including induction of astrocyte senescence. Aβ peptides, including soluble Aβ oligomers, correlate with disease severity [16] and upregulate p53, a key mediator of cellular senescence [17, 34]. Consistent with previous studies [16, 23], we demonstrate that Aβ increases p53 activity and its downstream targets, thereby inducing cellular senescence in primary human astrocytes (Fig. 1D-F). Aβ-induced astrocyte senescence is associated with upregulation of proinflammatory SASP cytokines (such as IL-6) and reduced expression of neurotrophic growth factors (such as IGF-1 and NGF) (Fig. 1G–1), which both contributed to astrocyte-mediated neurotoxicity in our previous studies on replicatively and radiation-induced senescent astrocytes [13, 14]. Our data in this study also indicate that the change in these soluble factors from Aβ-induced senescent astrocytes leads to neuronal apoptosis in trans-well co-culture experiment (Fig. 5, Vector control). Of note, in consistence with accumulation of DNA damage in AD brain tissues [13], our immunofluorescence staining of γH2AX and 53BP1 revealed that Aβ induces DNA double-stranded breaks in human astrocytes (Fig. 2), which cause p53 activation and senescence induction [30] possibly by a mechanism involving p38MAPK [16, 35]. The Aβ-induced DNA damage is likely due to oxidative stress-induced generation of DNA double-stranded breaks [28, 32, 36] and/or a direct nuclease activity of Aβ [31].

This study for the first time identifies Δ133p53α as an endogenous regulator of Aβ-induced astrocyte senescence and as a possible therapeutic target for Aβ-induced neurotoxicity (summarized in Fig. 6). Aβ diminishes the expression level of endogenous Δ133p53α in human astrocytes (Fig. 1B, C), and the lentiviral vector-mediated reconstitution of Δ133p53α expression prevents the cells from Aβ-induced DNA damage (Fig. 3), Aβ-induced senescence and SASP (Fig. 4) and Aβ-induced astrocytic neurotoxicity (Fig. 5). Although the effect of Δ133p53α on DNA damage (Fig. 3B, D), p21 expression (Fig. 4C) and IL-6 secretion (Fig. 4D) is significantly observed even without Aβ, it is much more evident against Aβ-induced changes. Our data suggest that Δ133p53α expression in astrocytes may be therapeutically enhanced in future applications to prevent or delay Aβ-induced pathological changes in AD.

**Fig. 6:**
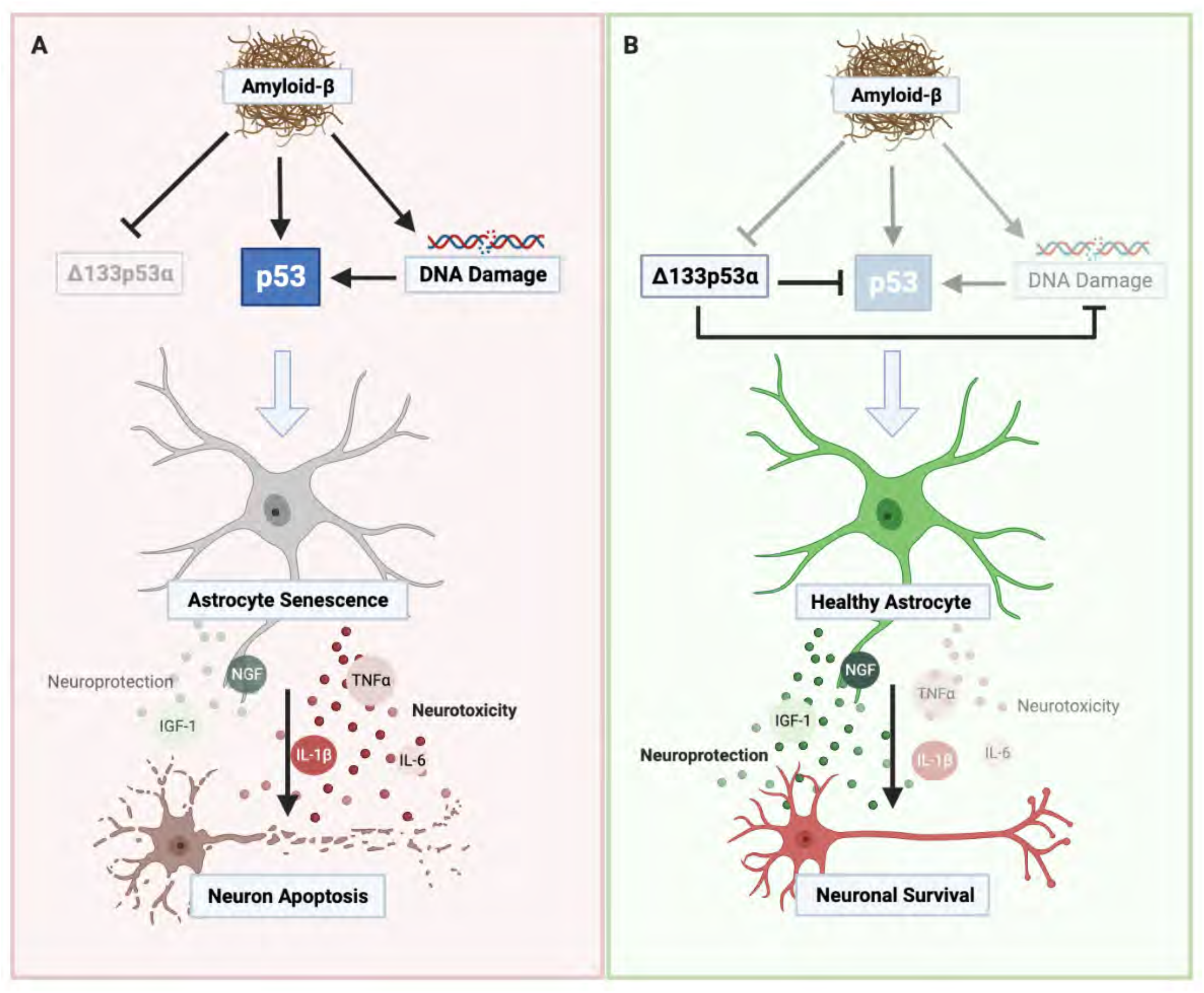
Δ133p53α in astrocytes and Aβ-induced neurotoxicity. **(A)** Aβ accelerates DNA damage and upregulates p53 in astrocytes without Δ133p53α-mediated inhibition, leading to astrocyte senescence and SASP. Astrocyte SASP not only promotes neuronal apoptosis but also generates and spreads proinflammatory environments, which in turn amplifies neurotoxicity. **(B)** Genetic enhancement of Δ133p53α (e.g., vector-driven expression in this study) or, ideally in future, its pharmacologic activation in astrocytes may constitute a therapeutic strategy for AD by disrupting such Aβ-triggered neurotoxic cascades.

Inhibition of p53 by expression of mutant p53 or use of a synthetic inhibitor has been shown to attenuate Aβ-mediated neuronal death and preserve cognitive functioning [37, 38]. However, such total inhibition of p53 could be mutagenic and oncogenic, thus not being considered for use in patients. In contrast, Δ133p53α has been shown to preferentially inhibit p53-mediated senescence while not inhibiting p53-mediated DNA repair or apoptotic elimination of damaged cells [20, 21, 39]. This selective dominant-negative inhibitory activity of Δ133p53α, together with its DNA repair-promoting activity [22, 33], well explains the beneficial effects of Δ133p53α observed in this study, and warrants further studies toward therapeutic applications.

## ABBREVIATIONS

(AD): Alzheimer’s disease
(Aβ): Amyloid-beta oligomers
(APP): Amyloid precursor protein
(NPCs): Neural progenitor cells
(SA-β-gal): Senescence-associated β-galactosidase
(SASP): Senescence-associated secretory phenotype

## ACKNOWLEDGEMENTS

This research was supported by the Intramural Research Program of the NIH and NCI-CCR. Dr. Beck is a Molecular Pathology Fellow in the NIH Comparative Biomedical Scientist Training Program supported by the National Cancer Institute in partnership with Purdue University. Fig. 6 was created with BioRender.com.

## AUTHOR CONTRIBUTIONS

KU, DL, CH designed the project. KU carried out experiments, data analysis and drafted the manuscript. All authors provided critical feedback and helped shape the research, analysis and manuscript. CH supervised the project.

## Supplementary Figures

**Supplementary Figure 1:**
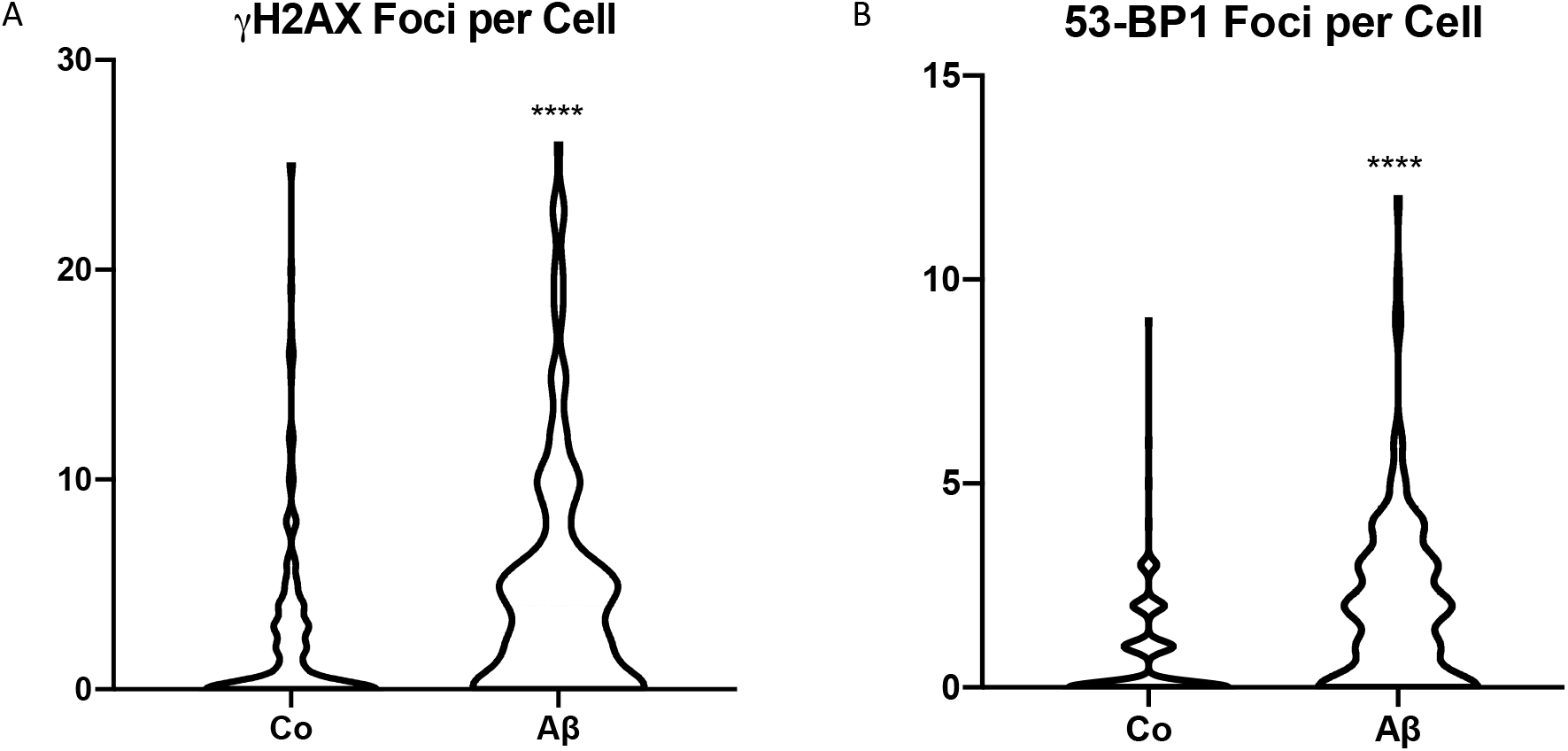
Amyloid-beta oligomers induce DNA damage in human astrocytes. Quantification of numbers of γH2AX foci (**A**) and 53BP1 foci (**B**) per cell 5 days after treatment with 1 μM Aβ for 24 hours. The pooled data from three independent experiments are shown as a single graph for each group. The same data are also presented as percentages of cells with 10 or more γH2AX foci (Fig. 2B) and numbers of 53BP1 foci per cell (Fig. 2C).

**Supplementary Figure 2:**
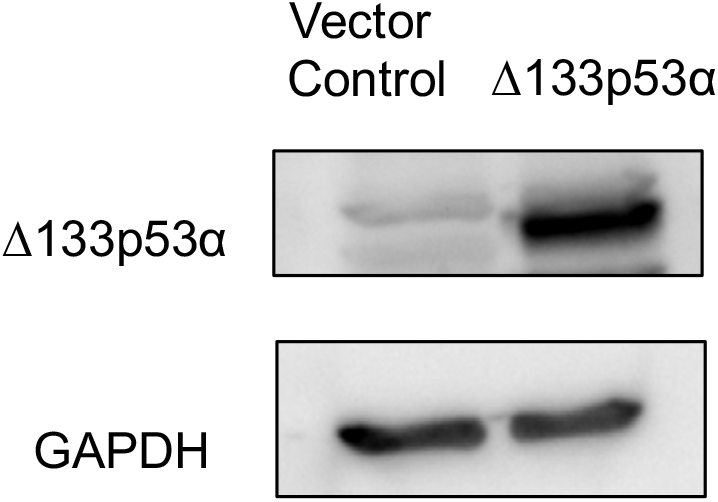
Confirmation of lentiviral vector-driven expression of Δ133p53α. Levels of Δ133p53α protein expression following transduction with the Δ133p53α lentiviral and control vector (pLenti6.3/TO/V5-DEST) were examined in immunoblot.

---

**Supplementary Table 1.**
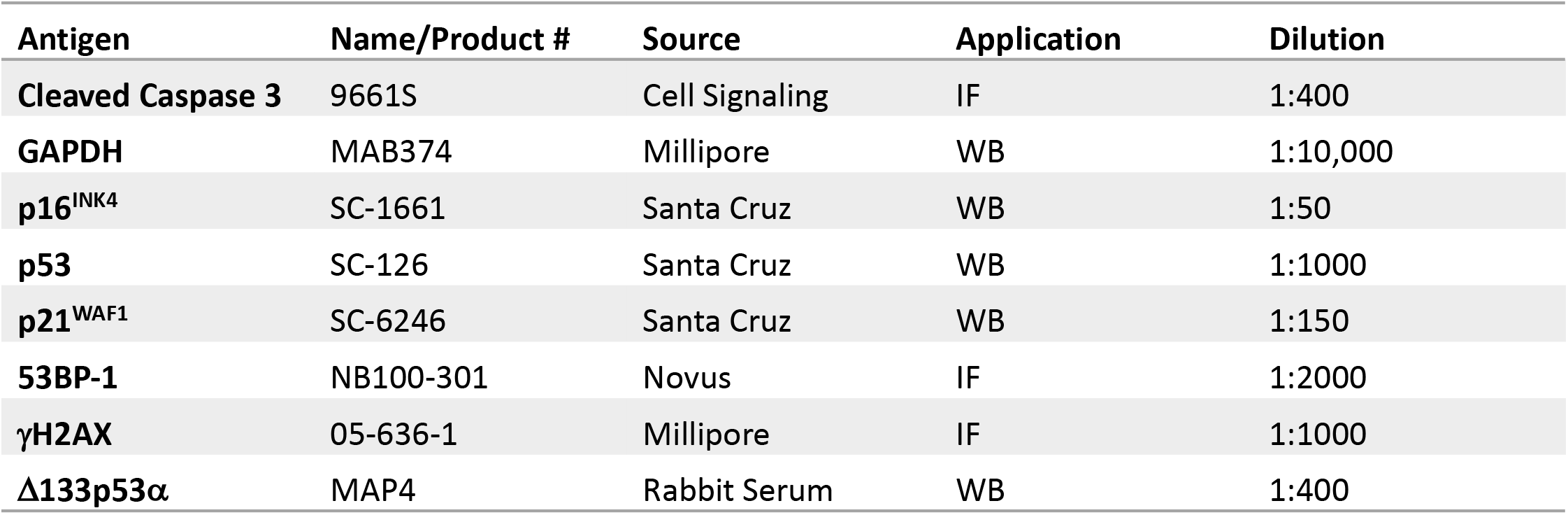

