## Supplementary Fig 1, 2 & Table for "“Δ133p53α Protects Human Astrocytes from Amyloid-Beta Induced Senescence and Neurotoxicity”"

#### Supplementary Figures

#### Supplementary Fig. 1

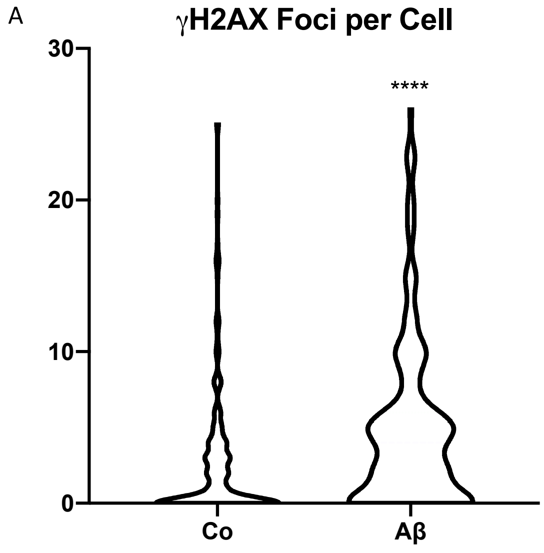


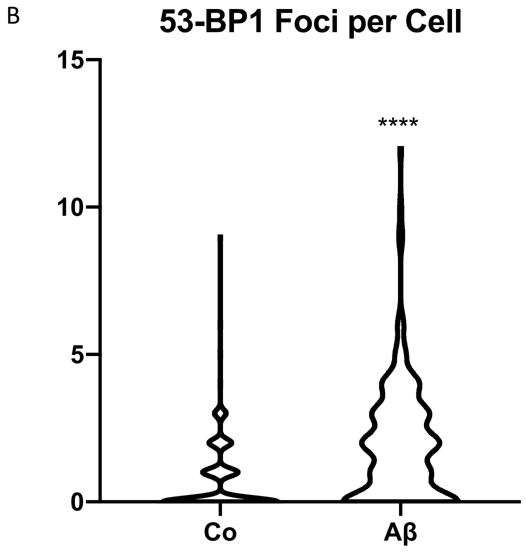


#### Supplementary Fig. 2

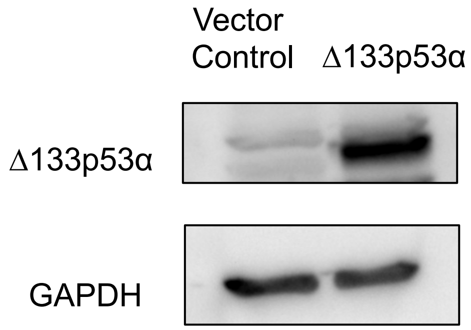

**Supplementary Figure 1: Amyloid-beta oligomers induce DNA damage in human astrocytes.** Quantification of numbers of γ-H2AX foci (**A**) and 53BP1 foci (**B**) per cell 5 days after treatment with 1 μM Aβ for 24 hours. The pooled data from three independent experiments are shown as a single graph for each group. The same data are also presented as percentages of cells with 10 or more γ-H2AX foci (Fig. 2B) and numbers of 53BP1 foci per cell (Fig. 2C).

**Supplementary Figure 2: Confirmation of lentiviral vector-driven expression of Δ133p53α.** Levels of Δ133p53α protein expression following transduction with the Δ133p53α lentiviral and control vector (pLenti6.3/TO/V5-DEST) were examined in immunoblot.

### Supplementary Table 1

| Antigen | Name/Product # | Source | Application | Dilution |
|---|---|---|---|---|
| **Cleaved Caspase 3** | 9661S | Cell Signaling | IF | 1:400 |
| **GAPDH** | MAB374 | Millipore | WB | 1:10,000 |
| **$p16^{INK4}$** | SC-1661 | Santa Cruz | WB | 1:50 |
| **p53** | SC-126 | Santa Cruz | WB | 1:1000 |
| **$p21^{WAF1}$** | SC-6246 | Santa Cruz | WB | 1:150 |
| **53BP-1** | NB100-301 | Novus | IF | 1:2000 |
| **γH2AX** | 05-636-1 | Millipore | IF | 1:1000 |
| **Δ133p53α** | MAP4 | Rabbit Serum | WB | 1:400 |
